# Non-Invasive Assays of Cochlear Synaptopathy -- Candidates and Considerations

**DOI:** 10.1101/565655

**Authors:** Hari M. Bharadwaj, Alexandra R. Mai, Jennifer M. Simpson, Inyong Choi, Michael G. Heinz, Barbara G. Shinn-Cunningham

## Abstract

Studies in multiple species, including in post-mortem human tissue, have shown that normal aging and/or acoustic overexposure can lead to a significant loss of afferent synapses innervating the cochlea. Hypothetically, this cochlear synaptopathy can lead to perceptual deficits in challenging environments and can contribute to central neural effects such as tinnitus. However, because cochlear synaptopathy can occur without any measurable changes in audiometric thresholds, synaptopathy can remain hidden from standard clinical diagnostics. To understand the perceptual sequelae of synaptopathy and to evaluate the efficacy of emerging therapies, sensitive and specific non-invasive measures at the individual patient level need to be established. Pioneering experiments in specific mice strains have helped identify many candidate assays. These include auditory brainstem responses, the middle-ear muscle reflex, envelope-following responses, and extended high-frequency audiograms. Unfortunately, because these non-invasive measures can be also affected by extraneous factors other than synaptopathy, their application and interpretation in humans is not straightforward. Here, we systematically examine six extraneous factors through a series of interrelated human experiments aimed at understanding their effects. Using strategies that may help mitigate the effects of such extraneous factors, we then show that these suprathreshold physiological assays exhibit across-individual correlations with each other indicative of contributions from a common physiological source consistent with cochlear synaptopathy. Finally, we discuss the application of these assays to two key outstanding questions, and discuss some barriers that still remain.

## 1. Introduction

Threshold audiometry is currently the foundation upon which clinical hearing evaluations are based. Accordingly, studies aimed at assessing the hearing damage associated with aging and acoustic overexposures have focused on *permanent* threshold changes between 250 and 8000 Hz (e.g., Rabinowitz et al., 2006; Cruickshanks et al., 2010). Temporary threshold shifts from noise exposure were considered relatively innocuous (National Institute of Occupational Safety and Health [NIOSH],1998). Human studies of age-related hearing loss tended to focus on individuals overs 60 years of age (e.g., Dubno et al., 2013). In contrast to these conventional views, animal data now show substantial permanent damage to synapses and auditory-nerve terminals innervating the cochlea (“synaptopathy”) from noise exposure that only causes temporary threshold shifts (Kujawa and Liberman, 2009; Lin et al., 2011; Gannouni et al., 2015; Bourien et al., 2014; Song et al., 2016), as well as with normal aging well before physiological changes characteristic of classic presbycusis (e.g., broad outer-hair-cell dysfunction) begin to manifest (Sergeyenko et al., 2013; Makary et al., 2011; Viana et al., 2015; Wu et al., 2018).

Unfortunately, from a clinical point of view, even an extreme degree of synaptopathy is unlikely to lead to changes in audiometric thresholds (Lobarinas et al., 2013; Furman et al., 2013). However, this “hidden” damage may have perceptual consequences (Bharadwaj et al., 2014; Plack et al., 2014; Schaette and McAlpine, 2011). Despite the common occurrence of potentially synaptopathic noise levels in everyday occupational and recreational settings, and emerging evidence of noise-induced synaptopathy in our non-human primate cousins (Valero et al., 2017), and in normally-aged human post-mortem tissue (Wu et al., 2018), the prevalence of cochlear synaptopathy in humans and its contributions to perceptual deficits remains unknown.

In order to understand the perceptual consequences of cochlear synaptopathy, it is essential to combine physiological measures of synaptopathy with perceptual measures in the same individuals. One strategy to achieve this would be to perform behavioral measurements in animal models in which synaptopathy can be directly assessed using microscopy and immunolabeling. However, it is possible that the behavioral consequences in relatively simple tasks are weak (e.g., see Oxenham, 2016) and that more complex listening conditions need to be created for the functional deficits to be apparent (Bharadwaj et al., 2014; Plack et al., 2014), rendering behavioral measurement in non-human animal models challenging. An alternate strategy, is to use non-invasive physiological assays that are putative correlates of synaptopathy in behaving humans and compare these measures to perceptual performance. Considerable effort is currently directed towards this enterprise by the hearing-research community.

The notion of comparing physiological correlates of processing in the early parts of the auditory pathway to auditory perception is not new. Indeed, otoacoustic emissions (OAEs), the auditory brainstem response (ABR), and the auditory steady-state response (ASSR), can each be used to estimate audiometric thresholds and detect clinical hearing loss (Gorga et al., 2003; Stapells & Oates, 1997; Lins et al., 1996). Studies comparing physiological measures to more complex perceptual tasks have typically relied on variants of the ASSR, such as the subcortical envelope-following response (EFR). For instance, EFR correlates of age-related declines in temporal processing (Fitzgibbons & Gordon-Salant, 2010; Snell & Frisina, 2000) have been reported in several studies (Purcell et al., 2004; Leigh-Paffenroth and Fowler, 2006; Grose et al., 2009; Ruggles et al., 2012). Even among young adults with normal audiometric thresholds in the clinical range, large variations in perceptual performance exist in challenging listening tasks (Kidd et al., 2007; Ruggles & Shinn-Cunningham, 2011). A portion of these individual differences in behavior correlate with both EFRs (Ruggles et al., 2011; Bharadwaj et al., 2015) and ABR measures designed to stress coding in the periphery (Mehraei et al., 2016; Liberman et al., 2016). Because these electrophysiological measures of subcortical coding are largely unaffected by top-down effects related to an individual’s state of arousal or attention (Varghese et al., 2015; Kuwada et al., 2002; Cohen & Britt, 1982; Thornton et al., 1989; See Section 3.6), these results suggest that that the fidelity of “bottom-up” neural processing very early along the auditory pathway can contribute to complex perceptual function. Finally, individuals experiencing tinnitus despite normal audiograms reportedly exhibit subtle differences in the ABR (Shaette & McAlpine, 2011), the EFR (Paul et al., 2017), and the acoustically evoked middle-ear muscle reflex (Wojtczak et al., 2017). Such results are consistent with the notion that cochlear synaptopathy contributes to important aspects of auditory coding in humans. However, whether that is truly the case is yet to be ascertained definitively.

Despite the many reports of correlations between aspects of perception and the integrity of neural processing in early parts of the auditory pathway, results exploring the association between risk factors for synaptopathy and non-invasive physiological measures such as the ABR are inconsistent. Studies comparing cohorts of young subjects with differing levels of acoustic exposure have reported ABR effects consistent with synaptopathy (Stamper & Johnson, 2015; Liberman et al., 2016; Skoe & Tufts, 2018; Bramhall et al., 2017). However, two larger studies that examined a wider age range found no association between estimates of noise exposure and the ABR or the EFR (Prendergast et al., 2017; Yeend et al., 2017). There are several possible explanations to these mixed results. It may be that humans are less susceptible to noise exposure than other species studied so far (Dobie & Humes, 2017), or that there are large variations in susceptibility across individuals such that exposure level *per se* in not a good predictor of damage (Davis et al., 2003). It is also possible that our ability to accurately estimate subjects’ acoustic exposure history is limited, given that one has to rely on individuals to report their past experiences (often many years to decades in the past). Another factor that can limit our ability to observe associations between non-invasive physiological measures and the degree of noise exposure or age is that these non-invasive measures can be affected by many sources of variability across individuals (and across repeated measurements within individuals) that are unrelated to synaptopathy. These extraneous factors that can affect non-invasive measures such as ABRs and EFRs are the subject of this report. First, we describe candidate non-invasive measures that can reflect synaptopathy and the evidence from animal models that motivate their use. We then systematically consider six extraneous sources of variability in these measures that can obscure the effects of synaptopathy. Finally, we show that with these extraneous factors carefully considered, three candidate synaptopathy measures exhibit across-individual correlations with each other; these correlations indicate contributions from a common underlying physiological source that is consistent with cochlear synaptopathy.

## 2. Candidate Measures of Cochlear Synaptopathy

### 2.1 Auditory Brainstem Response (ABR) Wave I Amplitude

Cochlear synaptopathy was first identified in CBA/CaJ mice following moderate noise exposure. This deafferentation was accompanied by only temporary elevations in distortion-product OAE (DPOAE) and ABR thresholds, but a permanent reduction in suprathreshold ABR wave I amplitudes (Kujawa & Liberman, 2009). The reduction in suprathreshold ABR amplitudes for tone-bursts of different frequencies correlated with the degree of synaptopathy found in the corresponding cochlear places, a result suggesting that suprathreshold ABR amplitude is a candidate non-invasive measure of synaptopathy. However, absolute ABR amplitudes do not appear reliable as a diagnostic in more genetically heterogenous animals. For instance, in a genetically heterogeneous cohort of guinea pigs with similar levels of synaptopathy as in the CBA/CaJ mice, absolute ABR amplitudes did not predict synaptopathic damage; only when suprathreshold ABR amplitude reductions (relative to pre-exposure amplitudes in the same ears) were computed were the ABR measurements related to synaptopathy (Lin et al., 2011; Furman et al., 2013). This suggests genetic heterogeneity can contribute variability to measures of absolute ABR wave I amplitude that is not easily normalized out in humans. In aging mice where immonolabeling showed cochlear synaptopathy, suprathreshold ABR wave I amplitudes were reduced in a manner similar to that found in noise-exposed mice. However, the relationship between synaptopathy and the ABR was most robust when the wave I amplitudes were normalized by the summating potential (SP; Sergeyenko et al., 2013). These observations suggest that some normalization procedure that reduces other sources of variability could be important when trying to interpret ABR measures.

In humans with tinnitus despite normal audiograms, Schaette & McAlpine (2011) reported that ABR wave I amplitude, normalized by wave V amplitude was reduced. This was interpreted as evidence of deafferentation at the auditory nerve level where wave I is thought to originate, and a compensatory “central gain” at the level of the midbrain where wave V is thought to originate. A similar result was found in mice with altered startle response properties following deafferentation, which was interpreted as a model for tinnitus (Hickox & Liberman, 2013). Along the same lines, in middle-aged rats, ABR wave I was reportedly reduced whereas wave V was relatively intact (Mohrle et al., 2016). These results suggest that wave V amplitude may be useful as a basis for normalization. However, as discussed in Section 3.3, the dominant contributions to ABR wave I and wave V might originate from different cochlear places when using broadband stimuli such as clicks, complicating the interpretation (Don & Eggermont, 1978).

The basic synaptopathic effects of noise exposure have now been observed in at least three species besides mice and guinea pigs -- chinchillas (Hickox et al., 2017), rats (Singer et al., 2013; Gannouni et al., 2016), and macaques (Valero et al., 2017). However, it is not yet established whether or not synaptopathy also manifests as robust reductions in suprathreshold ABR wave I amplitudes in these species.

### 2.2 Envelope-Following Response (EFR) Amplitude

Bharadwaj et al. (2014) hypothesized that cochlear synaptopathy, by virtue of being selective for nerve fibers with higher thresholds and lower spontaneous rates (Furman et al., 2013) may contribute to degraded EFRs at higher sounds levels and shallower modulation depths where high-spontaneous-rate nerve fibers tend to lose envelope timing (Joris & Yin, 1992). Consistent with this prediction, individual differences in human suprathreshold perception in complex tasks were correlated (∼25% of variance explained) with how robust one’s EFR amplitudes were to decrements in stimulus modulation depth (Bharadwaj et al., 2015). However, this hypothesis has not been directly tested in animal models with verified synaptopathy. Nonetheless, simpler EFR measures with high modulation rates (about 1000 Hz or more) have indeed been associated with synaptopathy in noise-exposed (Shaheen et al., 2015), and aging mice (Parthasarathy & Kujawa, 2018). EFRs at modulation rates beyond 800 Hz or so are thought to originate from the nerve by virtue of their short group delay (Shaheen et al., 2015) and based on the observation that that midbrain neurons do not temporally phase lock to modulations at those high rates (Joris et al., 2004). In this sense, unlike the typical human EFR experiments where modulation rates are lower (see Shinn-Cunningham et al., 2017 for a review), high-modulation rate EFRs are an alternate measure of nerve integrity much like the ABR wave I and may benefit from similar normalization procedures. For instance, Parthasarathy & Kujawa (2018) suggested that the high AM-rate EFRs that hypothetically originate from the nerve could be normalized by lower rate EFRs which may originate from the post-synaptic currents in midbrain neurons driven by input afferents; this is analogous to the wave I - wave V ratio.

### 2.3 Middle-ear Muscle Reflex (MEMR)

Perhaps the most promising non-invasive measure correlated with synaptopathy is the MEMR. The MEMR is typically measured acoustically as the change in middle-ear immittance properties induced by stimulus-driven efferent feedback to the middle-ear muscles. Single-neuron studies have raised the possibility that among afferent nerves, low-spontaneous rate (low-SR) nerve fibers dominate the input drive to the MEMR circuit (Liberman & Kiang, 1984; Kobler et al., 1992). Given the notion that the low-SR nerve population is more vulnerable to synaptopathy, this raises that possibility that the MEMR is a particularly sensitive candidate measure. Consistent with this, MEMR thresholds are elevated and suprathreshold amplitudes are attenuated in mice with synaptopathy, even when there is no hair-cell loss (Valero et al., 2017). Moreover, a strong correlation was observed across individual animals between the degree of synapse loss and MEMR thresholds (Valero et al., 2017). Another factor that might make the MEMR more sensitive than the ABR wave I is that low-SR contributions to the ABR appear to be modest (Bourien et al., 2014). Fortunately, from a clinical point of view, the MEMR can be measured rapidly, often in response to a single presentation of the eliciting stimulus. This facilitates the use of several different elicitors within a short period of time (e.g., noise bands with different center frequencies; See Valero et al., 2017). Moreover, MEMR measurement protocols are currently available on most clinical tympanometers and thus accessible to clinicians widely.

## 3. Extraneous Factors Modulating Non-invasive Assays of Synaptopathy

In this section, we discuss some of the extraneous sources of variability on the candidate synaptopathy assays described in Section 2. The extraneous factors are illustrated through the presentation of data from multiple experiments. All experimental data presented in this manuscript were acquired from adult (18 years or older) participants with clinically-normal audiograms, i.e., tone-based detection thresholds of 25 dB HL or better at standard audiometric frequencies up to 8 kHz. Data presented here were acquired at Boston University (OAE data in Section 3.3, ABR and EFR data in Section 4), Purdue University (ABR data in Sections 3.1 and 4, MEMR data in sections 3.5 and 4, acoustic calibration and audiometric data in Section 3.2), and Massachusetts General Hospital (EFR data in Section 3.4). All experiments were conducted using protocols approved by the local institutional review boards (IRBs). When relevant, additional participant details are provided alongside descriptions of each experiment in the following sections.

### 3.1 Basal Cochlear Gain Loss

The key finding with cochlear synaptopathy is that it occurs even in sections of the cochlea that do *not* show hair-cell loss (Kujawa & Liberman, 2015). However, concurrently with this cochlear synaptopathy in the mid sections of the cochlea, outer-hair cell (OHC) loss is sometimes seen in the far base (i.e., the “hook region”; Wang et al., 2002). Interestingly, in animal models where the hook-region OHCs and afferent synapses have both been examined, synapses in the mid and/or apical sections of the cochlea seem more vulnerable than the OHCs in the far base; hence, synaptopathy is almost always present in lower-frequency regions when high-frequency OHC damage is evident (Maison et al., 2013; Liberman et al., 2014). In humans, audiometric threshold elevation at “extended” high frequencies, i.e., at frequencies greater than 8 kHz, may be a reasonable proxy for such far basal OHC damage. Thus, elevated thresholds *beyond 8 kHz* could be a sign of cochlear synaptopathy *at lower frequencies* in humans. Indeed, the audiogram worsens continuously with age in humans with the loss progressing from high to low frequencies (Lee et al., 2012). Although the prevailing view is that noise-induced hearing loss first manifests as audiometric notches in the 4-6 kHz region, threshold elevation at extended high-frequencies has also been reported in young humans with above average noise exposures with clinically normal thresholds up to 8 kHz (Liberman et al., 2016), and has been suggested as an early marker for noise-induced damage (Mehrparvar et al., 2011).

We examined the relationship between audiometric thresholds beyond 8 kHz (averaged over 10, 12.5, 14 and 16 kHz) and ABR wave-I amplitudes evoked by 80 dB nHL clicks recorded using clinical equipment at Purdue University (Figure 1). As seen in Figure 1, higher average thresholds in the 10-16 kHz range is significantly associated with lower wave I amplitudes consistent with the idea that cochlear synaptopathy co-occurs with damage to OHCs in the far base of the cochlea (Pearson r = −0.44, N = 136, p=9e-10). Moreover, an interesting feature of the data in Figure 1 is that it shows a “lower-triangular” pattern as highlighted by the dotted triangle; that is, we see many cases of small ABR wave-I values despite good thresholds beyond 8 kHz, but the other way around is much less common. The lack of data in the the top-right corner in the scatter is statistically significant (p = 0.0003) based on the non-parametric test described by Bardsley et al., (1999). These data are consistent with the idea that one could have synaptopathy without OHC damage in the far base, but when OHC damage is present, broader synaptopathy is almost always concomitant.

**Figure 1:**
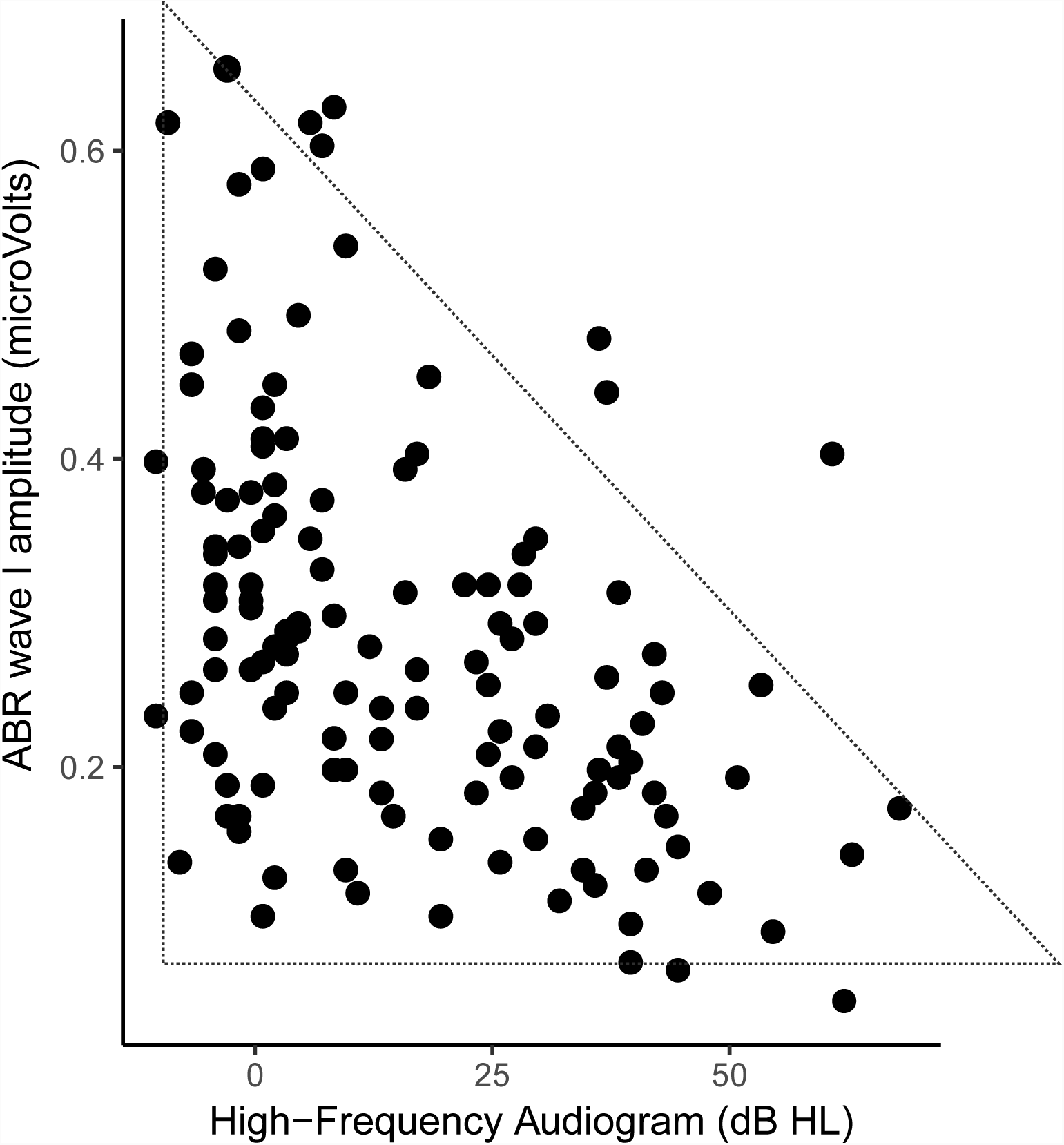
Relationship between ABR wave I amplitudes and extended-high-frequency audiometric thresholds (averaged over 10 – 16 kHz) for 136 ears with clinically normal thresholds (better than 25 dB HL) up to 8 kHz. Greater thresholds in the 9-16 kHz range are associated with smaller wave I amplitudes. Moreover, a lower-triangular pattern of scatter is evident, i.e., there are many ears with small wave I amplitudes despite good 10-16 kHz thresholds, but very few data point with the opposite trend. These results are consistent with the interpretation that when there is OHC damage in the far basal parts of the cochlea, broader cochlear synaptopathy is also present. An alternate interpretation is that the 9-16 kHz region of the cochlea is a prominent contributor to the wave I.

Although this interpretation is tempting, an alternate explanation must be considered. Don & Eggermont (1978) showed that the ABR wave I reflects contributions from the base of cochlea. Even if the click stimuli are band limited (here ER-3A insert earphones were used that roll-off starting at around 4 kHz), upward spread of excitation could recruit contributions from sections of the cochlea that are tuned to 10-16 kHz. Thus, the observed correlations could just indicate that cochlear gain loss in the 10-16 kHz region reduces the contribution of this region to the ABR wave-I, thereby reducing the wave I amplitude. The “lower-triangular” pattern, however, is not as easily explained with this interpretation.

These two possible competing interpretations also arise in the context of correlations between speech-in-noise perception and extended high-frequency audiograms. There is some evidence, although sparse, that individuals complaining of speech-in-noise problems despite normal audiograms show elevated thresholds beyond 8 kHz (Badri et al., 2011; Shaw et al., 1996). This raises the question of whether audibility in those frequencies is intrinsically important for speech-in-noise perception, or whether threshold elevation at those frequencies is a marker for other damage, including cochlear synaptopathy at lower frequencies.

To disambiguate between the two competing interpretations for the correlation between the ABR and extended-high-frequency audiograms, the most direct test would be to compare the ABR amplitudes in animals with and without OHC loss in the hook region, and with and without broader cochlear synaptopathy, if at all such selective damage is achievable. The correct interpretation of the correlations is likely a combination of both of these views, with no strong evidence yet to support one view more than the other. In either of those cases, however, this extraneous factor of high-frequency OHC loss should be considered. One approach to circumnavigate this issue for the purpose of assaying cochlear synaptopathy would be to “regress out” (or otherwise statistically account for) the audiometric variations beyond 8 kHz from ABR measures. Any residual relationship between the ABR and risk factors such as noise-exposure and age can then be reasonably attributed to mechanisms distinct from OHC damage. However, this approach is likely too conservative because cochlear synaptopathy is correlated with OHC damage, and regressing out audiograms might attenuate the effects attributed to syanptopathy. This might contribute to an elevated rate of false negatives, i.e., a bias towards reporting a lack of correlation between ABR and noise exposure, or ABR and age. Nonetheless, measuring audiograms beyond 8 kHz would be useful in studies involving any cohorts of human subjects that are at risk for synaptopathy.

### 3.2 Ear-canal Effects

It is well known that the acoustic pressure and intensity of stimuli delivered to the ear can depend on the immittance properties of the outer and the middle-ear. Accordingly, calibration procedures of supra-aural, circum-aural, and insert earphones in hearing science and clinical audiology tend to use cavities that mimic the average human ear as established in International Standards (IEC-60318; International Electrotechnical Commission). Calibrations using such standards, while adequate for the purposes for which they were designed, do not account for individual variations in acoustic properties of the ear. With insert earphones, the effects of the ear canal properties can also depend on the placement of the transducer couplers in the ear canal (e.g., shallow vs. deep insertion), an effect that is most evident at higher frequencies (Siegel, 1994). Souza et al., (2014) showed that moving the insert coupler by as little as a few mm in the ear canal can produce up to a 20 dB change in audiometric thresholds at frequencies greater than 3 kHz. Although lower-impedance circumaural headphones should theoretically be less affected by the properties of the ear, such filtering effects can significantly limit the interpretability of extended-high-frequency audiograms such as those discussed in Section 3.1. A primary contributor to this is the standing wave interference pattern from back-and-forth reflections of sound that occur between the tympanic membrane and the insert coupler. These effects are well described in the OAE/immittance literature with many compensating strategies or alternate calibrations proposed (Scheperle et al., 2008; Souza et al., 2014; Charaziak et al., 2017). Of these, forward-pressure-level (FPL)-based calibrations have been shown to be robust and have the advantage of being able to control the phase of stimulation that drives the middle-ear at the tympanic membrane (Souza et al., 2014). FPL is the resultant level of the superposition of all forward traveling wavefronts that arise from repeated reflections in the ear canal space. Fortunately, FPL can be estimated accurately using a microphone in the ear canal (as available with OAE probes) and *a priori* sound source calibrations using classic analysis techniques developed for two-port systems (e.g., see Keefe et al., 1992).

Although ear-canal insertion depth for a given listener has been emphasized in the OAE literature, the variability introduced *across individuals* for a nominal insertion has not been systematically studied, to our knowledge. Here we examine this question in two ways: (1) we compare the estimated forward-pressure levels (FPL; Scheperle et al., 2008) across listeners for a fixed voltage applied to the speakers and for a nominal insertion that is representative of typical experiments, and (2) we compare audiometric thresholds in the 8-16 kHz range obtained for a cohort of individuals using a standard SPL calibration (tested using circumaural headphones) against FPL-based audiograms obtained using the ER-10X OAE probe (Iseberg et al., 2015) for the same cohort of subjects.

Figure 2 shows the voltage-to-FPL transfer functions obtained on three different individual listeners for a relatively deep insertion of the ER-10X probe for each subject. It is evident that with typical setups, there is about a 15 dB variation at some higher frequencies across individuals. This shows that when one is using insert earphones with SPL-based calibrations, individual variations in ear anatomy can introduce considerable level variations in the forward-traveling pressure wave at the tympanic membrane. Given that the energy transmitted through to the inner ear is likely to be more closely related to physiological and perceptual responses than energy that would be incident from a speaker for a given voltage, this can be a significant source of variability in any narrow-band measurements. Indeed, for data from 88 ears, we find that FPL-based audiograms in the 8-16 kHz range tend to be monotonic (72 ears, i.e., 82% of the ears). In contrast, even when using circumaural earphones, audiograms based on SPL-calibration (in a standard cavity) exhibited a greater rate of non-monotonicity (i.e., idiosyncratic peaks or valleys; only 59 ears, or 67% of ears showing a monotonic audiogram). Note that non-monotonicity was defined as any increase then decrease (or decrease then increase) in audiometric thresholds between adjacent frequencies in the 8 kHz to 16 kHz range (i.e, at 8, 10, 12,5, 14, and 16 kHz). Also note that all audiogram data were expressed relative to the mean subject’s thresholds (i.e., in dB HL units) for this comparison, and that the resolution for thresholds was 5 dB. Using the FPL-calibrated audiograms’ rate of non-monotonicity as the reference data for a binomial test, the standard audiogram’s rate of non-monotonicity is significantly higher (p = 0.0005), likely reflecting ear-canal filtering effects. Thus, when frequency-specific assessment is desired, we recommend that FPL-based calibrations be employed for assays of cochlear synaptopathy.

**Figure 2.**
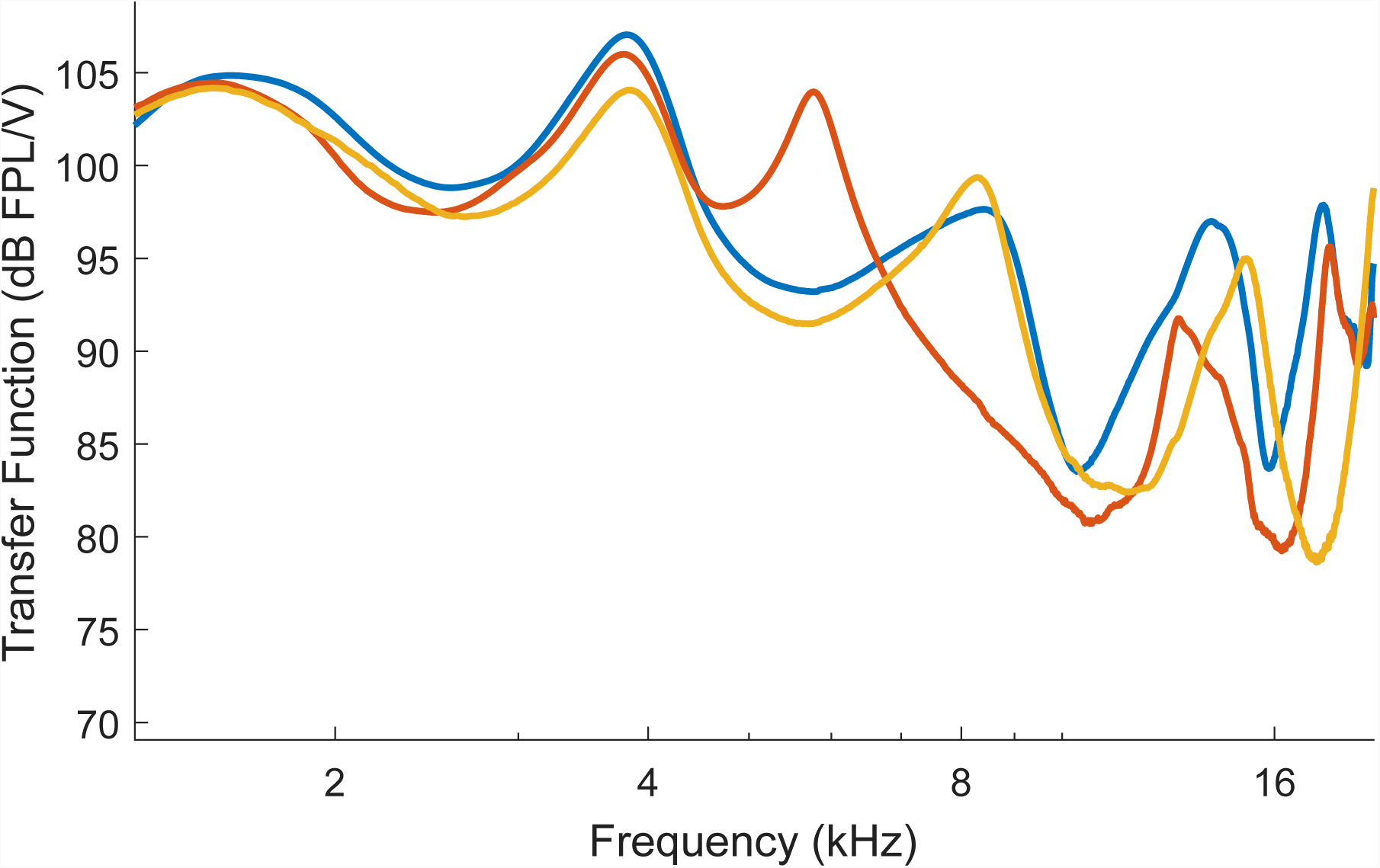
Voltage to forward-pressure-level (FPL) transfer functions obtained from three different ears using the ER-10X OAE probe for typical probe insertion depths. Considerable variability is seen in the FPL levels for a constant voltage input across ears at higher frequencies. This suggests that when conventional calibration techniques are employed, individual differences in ear-canal filtering could contribute to variability in the stimulus driving the middle-ear for insert probes.

### 3.3 Cochlear Mechanical Dispersion

Assays of suprathreshold hearing in humans, by virtue of being non-invasive, reflect population responses along the auditory pathway. Thus, in addition to the response properties of single-neurons affecting such measures, the relationship *between* the responses of the thousands of neurons in the population likely matters. For electrophysiological responses such as ABRs and EFRs, this multisource population activity produces scalp potentials that depend on (1) how synchronous the responses are across the individual neural currents that make up the overall response, and (2) the geometry of the source currents relative to the recording electrodes, and the conductivity profile of the tissue volume in between (Hubbard et al., 1971, Okada et al., 1997, Irimia et al., 2013). The latter effects will be discussed in Section 3.4. Here, we focus on the synchrony of responses across different neural currents.

One factor that can affect the synchrony of responses across the auditory-nerve population is the level-dependent cochlear traveling wave delay, with the base being excited before the apical half (Shera & Guinan Jr., 2003). Both ABRs and EFRs can be affected by systematic individual differences in the anatomy and mechanics of the cochleae that lead to these dispersive effects (Don et al., 1994; Nuttall et al., 2015). This dispersive effect is thought to underlie the sex differences observed in ABR amplitudes (Don et al., 1993). Indeed, a human female cochlea is about 13% shorter on average than a male cochlea, but with a similar range of frequencies represented along the tonotopic map, indicating a 13% larger stiffness gradient (Sato et al., 1991). This could translate to faster base-to-apex response propagation times within the cochlea of females compared to males, which in turn could lead to more synchronized responses from different portions of the cochlea, producing larger amplitudes for the measured population response. Consistent with cochlear dispersion being an important factor affecting ABRs, computational models that incorporate different cochlear mechanical models produce considerably different predictions for ABR amplitudes and latencies and how they vary with level (Verhulst et al., 2015). Here we illustrate this important source of variability by showing that click-evoked OAE group delays are shorter for female than male subjects consistent with the ABR wave I being larger for female ears (Don et al., 1993).

Figure 3A shows the group delay obtained from broadband-click-evoked OAEs for frequencies around 2 kHz. Consistent with the idea that mechanical response propagates slower on average in male subjects, the group delay is slightly longer in males than females. Note that the OAE group delay consists of the delay from not only the forward propagation and filter buildup time, but also the reverse propagation of the reflection emission. This suggests that some normalization procedure on the amplitude could be useful. At a minimum, analyses must take into account sex effects on the ABR. One candidate for a normalization denominator is the ABR wave V because it is seemingly less affected than wave I by deaffarentation of the periphery (see Section 2.1). Unfortunately, however, the ABR wave I and wave V are thought to arise from overlapping but different sections of the cochlea (Don & Eggermont, 1978). Here we illustrate this issue by comparing the relative amplitudes of wave I and wave V (i.e., the I/V amplitude ratio) for conventional broadband clicks and clicks that are high-pass filtered at 3 kHz. Stimuli in both cases were delivered at 80 dB above the detection thresholds for three pilot subjects. The broadband click level was comparable to standard 80 dB nHL clicks in intensity. Figure 3B shows the ABR wave I-to-wave V ratios obtained from the same subjects for broadband and high-pass clicks. Clearly, the absolute value of the ratio is altered such that wave I is larger than the wave V for high-pass clicks and wave V is larger than the wave I for broadband clicks (on average). This is consistent with the notion that cochlear contributions for wave I and wave V are not the same, with wave I emphasizing more high-frequency cochlear sections than wave V, as previously reported (Don & Eggermont, 1978).

**Figure 3.**
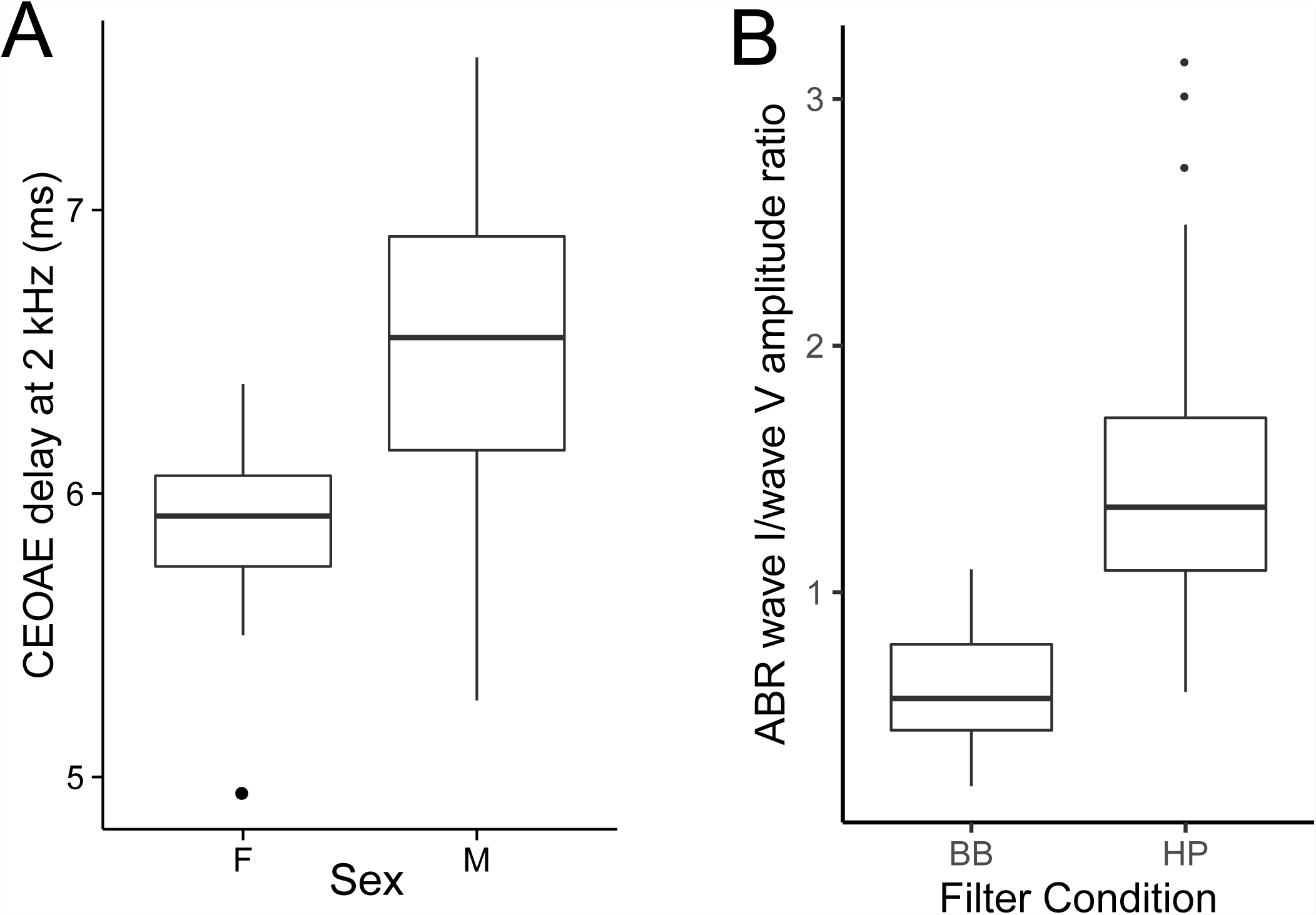
OAE and ABR data illustrating the dispersive mechanics of cochlear excitation. Panel A shows click-evoked OAE group delays for frequencies around 2 kHz (corresponding to roughly the middle of the cochlear spiral). Consistent with larger ABR amplitudes seen in female subjects, OAE group delays are shorter indicating that the dispersive effects of the cochlear traveling wave are less pronounced in female ears. Individual differences in such cochlear dispersion can contribute to variability in ABR amplitudes. Panel B show ABR wave I to wave V amplitude ratios obtained for broadband (BB) and high-pass (HP) click of approximately the same sensation level. For BB clicks, the wave V is larger consistent with a greater contribution of low-frequency portions of the cochlea to wave V compared to wave I. In contrast, with HP clicks, the relationship is reversed (as with tone-pip data in animal models). These observations suggest that when considering amplitude ratios, it is important to account for differences between the cochlear regions recruited by wave I and wave V.

Overall, these results suggest that some normalizing procedure for ABR wave I amplitudes could be beneficial in reducing the dispersive effects of cochlear response times, but also that when broadband clicks are used, the wave I-to-wave V ratio is not easily interpreted. Thus, one possibility is to use high-pass clicks and apply the normalization procedure, as we do in Section 4. Another viable candidate for the normalization denominator is the hair-cell summating potential (SP), as illustrated in Sergeyenko et al. (2013) and Liberman et al. (2016). In our lab, the SP is consistently observed only for click levels of ∼110 dB peSPL when using ear-canal electrodes (tiptrodes) and may be harder to obtain in clinical settings. When obtained, the SP could be used for normalization. Because the SP is generated presynaptically, the interpretation is more straightforward than when using wave V as a reference. Yet, how cochlear mechanical dispersion is manifested in the amplitude of the SP is currently unknown.

### 3.4 Volume Conduction Effects

As mentioned in Section 3.3, both the geometry of the neural source currents relative to the recording electrodes and the geometry and the conductivity of the intervening tissue volume can affect scalp-measured voltage responses (Hubbard et al., 1971, Okada et al., 1997, Irimia et al., 2013). A consequence of this fact is that individual anatomical variations and variations in electrode positioning can both introduce undesirable variations in these non-invasive electrophysiological measures.

To examine the contribution of individual variations in anatomy, we compared individual differences in EFRs measured using electroencephalographic (EEG) and magnetoencephalographic (MEG) data. The idea behind this strategy is that MEG and EEG have markedly different sensitivity profiles to source currents in different parts of the brain, and are affected differently by the tissue volume and boundaries (Hamalainen et al., 1993). Indeed, the physics of MEG and EEG recordings dictates that EEG is more sensitive to neural currents that are oriented radially to the scalp surface whereas MEG is more sensitive to tangential sources (Hamalainen et al., 1993; Ahlfors et al., 2010). Thus, if MEG and EEG measures of EFR provided similar ranking of individuals based on the respective EFR amplitudes, that would suggest only a small contribution from anatomical factors and electrode positioning. On the other hand, if the ranking are inconsistent across the two measures, that would suggest a significant contribution from anatomical and electrode-positioning factors.

Crucial to this direct comparison of MEG-and EEG-based ranking of individuals is the assumption that MEG and EEG measures are picking up a common underlying neural source and are different only in how sensitive different sensors are to this common source. If this assumption is satisfied, then the differences in ranking of subjects from MEG and EEG should come primarily from anatomical factors, electrode positioning, and measurement noise. To test this assumption, we first examined the EFR phase and group delay for MEG and EEG measurements (see Shinn-Cunningham et al., 2017 for a discussion). Figure 4A shows the EFR phase response for different envelope frequencies imposed on a broadband noise for MEG and EEG, respectively. Interestingly, the estimated group delay is more than twice as large (∼19 ms) for MEG around 80-100 Hz than for EEG (∼8 ms). This suggests that EEG-based EFRs near 100 Hz (a commonly used frequency for EFR measures in humans) weight sources earlier along the auditory pathway more strongly, and MEG-based EFRs near 100 Hz have greater contributions from hierarchically later sources. This result is consistent with older reports showing disparities in the group delay and source localization between EEG-based and MEG-based auditory steady-state responses (Schoonhoven et al., 2003; Ross et al., 2000; Herdman et al., 2002), and with recent source-localization and group-delay data showing that EEG-based EFRs above 80 Hz or so are dominated by subcortical sources (Bidelman et al., 2018; Shinn-Cunningham et al., 2017), whereas MEG-based measures could have some cortical contribution (Coffey et al., 2016). Unlike around 100 Hz, EFRs at frequencies above 200 Hz showed a similar group delay with MEG and EEG, suggesting a common pattern of sources dominated by subcortical nuclei. Consistent with this notion, MEG gradiometers, which are less sensitive to deep sources than magnetometers, did not show an EFR at 223 Hz (Figure 4B). Thus, for examining the contribution of individual differences in anatomy to the EFR, we compared MEG and EEG responses at an envelope frequency of 223 Hz.

**Figure 4.**
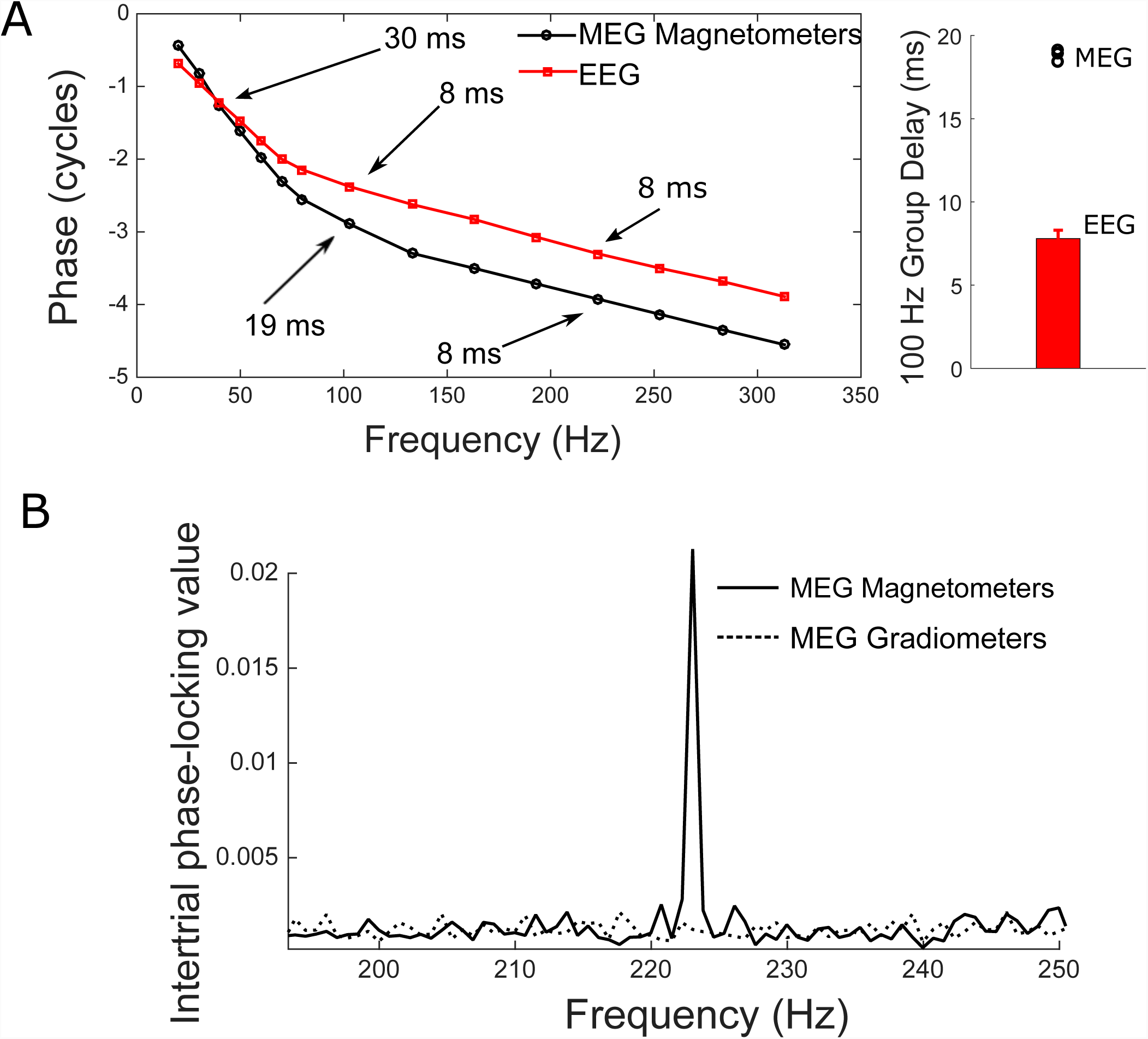
Comparison of MEG and EEG versions of EFR measures. Panel A (left) shows the response phase vs. frequency functions for EFRs obtained from MEG and EEG for a representative subject. The differences in slope near 100 Hz, and the similar slopes beyond 200 Hz are evident. The group delays extracted from the phase for three MEG subjects are shown along with EEG group delays estimated from 10 subjects (Panel A, right). For the 100 Hz EFR, MEG group delay is more than twice as long as EEG suggesting that MEG and EEG versions of EFR can only be compared for modulation frequencies beyond 200 Hz or so. Panel B shows the EFR response at 223 Hz for two types of MEG sensors. The gradiometers which are insensitive to farther sources do not show a response, whereas magnetometers do. This is consistent with a subcortical source dominating the MEG response for this modulation frequency.

We ranked eight adult subjects from 1 through 8 based on the MEG EFR amplitudes for the best channel for each subject (this coincided with the ranking that would be obtained using source amplitudes with an equivalent-current dipole fit approach; Table 1 top row). We then ranked the same eight subjects using EEG-based EFR amplitudes at the Cz scalp location relative to the earlobes. This was done to mimic clinically-viable EFR recordings with electrodes placed at nominally the best locations for single channel recordings (i.e., Cz and earlobes). The rankings obtained are shown in Table 1 (middle row) and correspond to a rank correlation (Kendall tau) of 71% (p = 0.01) with MEG rankings. This suggests that although MEG and EEG-based EFR measures are significantly correlated, there is some scrambling of ranks between measures, likely from anatomical factors and electrode positioning. We also ranked the same subjects based on multichannel EFR amplitudes obtained using the complex-spectral principal component analysis (cPCA) method described in Bharadwaj & Shinn-Cunningham (2014). The multichannel EEG-based rankings are shown in Table 1 (bottom row) and correspond to a rank correlation of 85% (p = 0.002) with MEG rankings. Note that rank correlations rather than Pearson correlations are reported here because test-retest rank correlation of absolute EFR amplitudes (i.e., measures on the same individuals in two separate sessions with EEG) tend to be 100%, whereas test-retest Pearson correlations are lower. The MEG-EEG comparisons suggest that combining multiple EEG channel using the cPCA method can reduce the effect of anatomical factors and electrode positioning by giving a more stable estimate of the EFR response. Overall, these results suggest that anatomical factors do contribute to individual differences in EFR amplitudes. One may expect similar results for ABR wave V amplitudes. Thus, for both ABRs and EFRs, using multichannel recordings could help mitigate variability arising from anatomical differences and electrode positioning variations.

**Table 1.** Ranking of individual subjects based on EFR amplitudes obtained from MEG or EEG

|  |  |
| --- | --- |
| MEG-based ranking of subjects (reference ranking) | 1, 2, 3, 4, 5, 6, 7, 8 |
| EEG-based ranking of subjects from single channel (Cz). Subjects are listed in rank order from reference ranking. | 1, 3, 2, 6, 5, 4, 7, 8 |
| EEG-based ranking of subjects from multichannel complex PCA method. Subjects are listed in rank order from reference ranking. | 1, 2, 3, 5, 6, 4, 7, 8 |

### 3.5 Immittance and Reflex Spectra

Typical clinical MEMR measurements are performed acoustically using a pure tone probe at 226 Hz. However, the middle-ear is a broadband transducer with stereotypical immittance spectra (Feeney et al., 2017). Accordingly, the acoustically evoked MEMR is also a wideband change characterized by a reduction in the absorbed power at low frequencies along with alternate bands of increases and decreases at various higher frequencies (Keefe et al., 2017). Crucially from the perspective of using the MEMR as a measure of synaptopathy, there could be variations in the profile of the MEMR with frequency across individuals. This source of variability is undesirable; for example, when using the classic clinical measure of MEMR, an individual whose MEMR spectrum happens to peak near 226 Hz might artificially appear to have lower thresholds and larger MEMR amplitudes compared to another individual whose MEMR spectrum peaks farther from 226 Hz. This spectral variance can be even more exaggerated when using raw ear-canal pressure changes to measure the MEMR rather than using probe calibrations to measure the reflex as a change in absorbance or absorbed power. Here, we illustrate these issues by (1) comparing wideband measurements of the MEMR using a click probe to tone-based measurements in the same individual, and (2) by comparing the spectra of ear-canal pressure change induced by an MEMR-eliciting stimulus across individuals.

Figure 5A shows the MEMR measured on an individual subject using a wideband probe (a click) or a series of pure tones (200 to 1600 Hz) to mimic measurement done with standard clinical protocols but over a wider range of frequencies. For this experiment, the MEMR is quantified as the change in ear-canal pressure induced by a 76 dB SPL broadband noise elicitor. Wideband measures and tone-based measures were interleaved and the pressure changes measured for tonal probes were scaled up to account for the difference in spectral level between the click at a given frequency and the individual tone at that frequency. As seen in Figure 5A, the tone-based measures simply are frequency samples of the wideband measure, as expected from the general linearity of acoustic measurements. However, the wideband measure using a click probe is significantly more efficient in obtaining the same information. Figure 5B shows wideband MEMR spectra (quantified as ear-canal pressure change again) for two different individuals as a function of elicitor level. Although the subject on the right panel in Figure 5B has a higher threshold (around 76 dB FPL) and overall smaller MEMR amplitudes compared to the subject on the left panel (around 52 dB FPL), the MEMR spectra are different in shape, with the subject on the left panel showing a relatively small change around 226 Hz (frequency used in typical clinical MEMR measurements) with the other subject showing larger changes. This suggests that single frequency measurements of the MEMR could be problematic when comparing across individuals. While the spectra do become more stereotypical when using absorbed power changes or absorbance changes to quantify the MEMR (Keefe et al., 2017), the low-frequency end shows considerable variability in spectral shape across subjects and is also noisier. Indeed, Feeney et al. (2017) showed that wideband measurements allow for detection of an MEMR at lower elicitor levels compared to classic 226 Hz tone-based measurement, suggesting greater resistance to measurement noise and spectral profile variations. These observations suggest that wideband measurements should be preferred when using the MEMR as an assay of synaptopathy where the MEMR spectra can be summarized by averaging over a broader range of frequencies to reduce the effects of individual spectral-profile peculiarities. Wideband immittance and reflex measurement protocols are starting to be available in clinical tympanometers and likely will be accessible to interested clinicians more widely in the near future.

**Figure 5.**
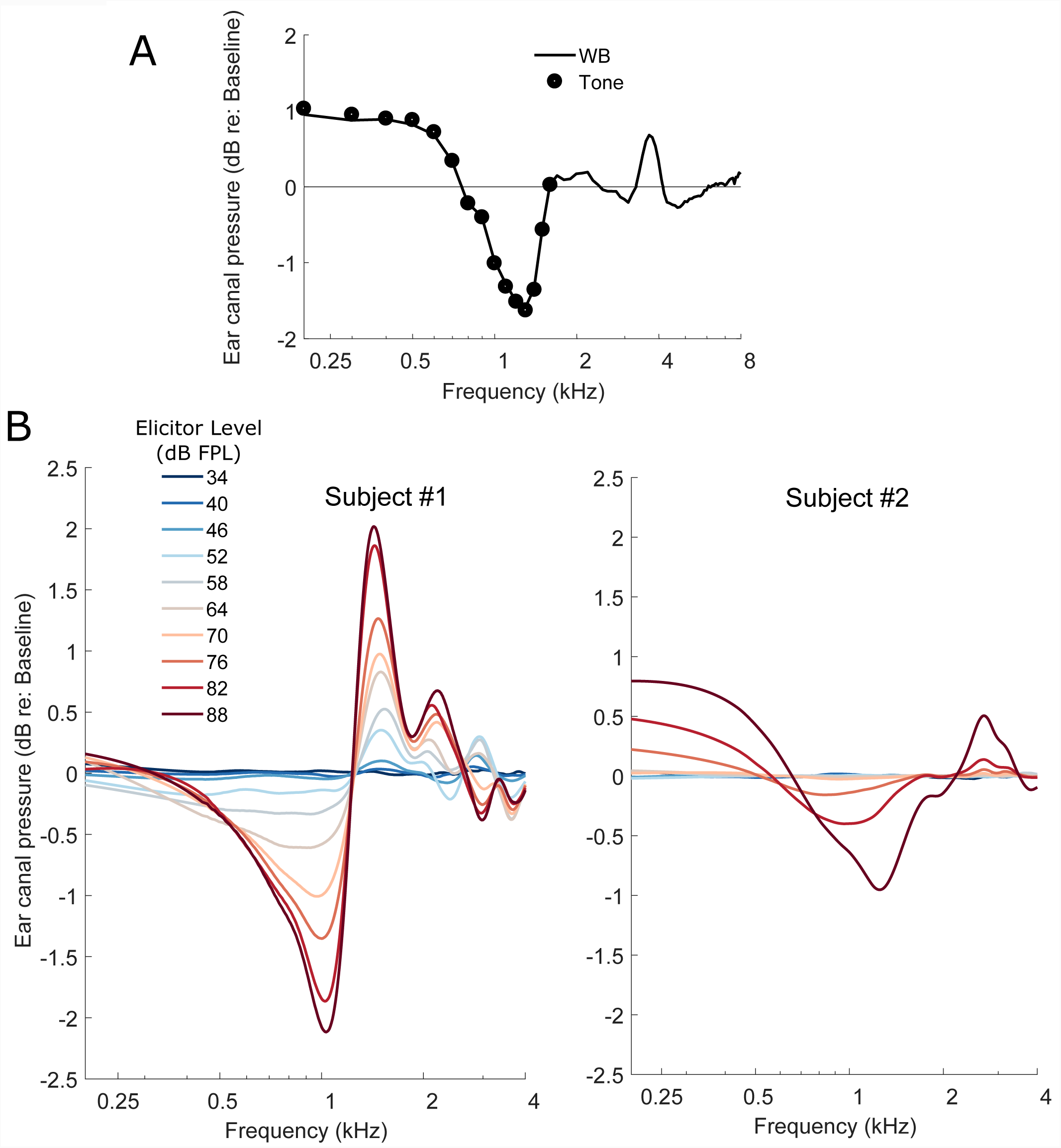
Data illustrating the spectral profile variability of the MEMR. Panel A shows the MEMR elicited by a 76 dB FPL broadband noise for an individual. The MEMR was measured using either a click probe, i.e., a wideband (WB) measurement, or using tone probes in interleaved trials. The coincidence of the WB spectrum with the individual data points from different tone probes confirms the linearity of acoustic measurement and suggests that a WB measurement is much more efficient. Panel B shows the MEMR spectra obtained for two different subjects (left and right) for a series of broadband noise elicitors at different forward-pressure levels. Both subjects show an increasing MEMR response as the elicitor level is increased. However, the spectral shapes are different; the individual in the left panel shows small changes near 226 Hz although a large response at other frequencies. In contrast the individual shown on the right has small responses overall, but shows larger changes near 226 Hz. Thus, if the MEMR were to be measured only at one frequency (say 226 Hz), the ordering of who has a larger response would be swapped.

### 3.6 Effects of Arousal and Attention

It is sometimes thought that the degree of arousal (e.g., awake vs. asleep) or selective attention to a target sound that is eliciting the ABR or EFR can modulate those responses. This is thought to be possible either through corticofugal feedback or through efferent control of the auditory periphery. Here, we wish to draw a distinction between effects of attention and arousal that may be present and measurable through detailed physiological recordings, and effects on non-invasive assays such as ABRs and EFRs per se, which reflect the aggregate response of thousands of single neurons. The vast majority of ABR and EFR studies examining the effect of attention have reported null results or reported effects that are very small compared to the range of individual differences (See Varghese et al., 2015 for a discussion). Indeed, in clinical settings, ABRs are routinely recorded under sedation and interpreted in the same way as awake ABRs. When (presumably) subcortical EFRs and cortical responses were measured simultaneously, the EFRs show no effects of attention even when strong effects are seen at the cortical level (Varghese et al., 2015). We informally analyzed the EFR amplitudes of four subjects who fell asleep halfway through an EFR recording. Because we recorded EFRs with a 32-channel EEG cap, we were able to use the low-frequency portion of the EEG to reliably extract 600 trials during stage-2 or slow-wave sleep and compare them to 600 trials where they were awake. The magnitudes were indistinguishable from each other with an across-subject Pearson correlation of 0.98. The noise floor, however, changes considerably (∼12 dB for one subject around 100 Hz). Similar observations have been made in the OAE literature, where an attention task can affect the noise floor without affecting the actual evoked OAE (Francis et al., 2018).

The fact that ABRs and subcortical EFRs are relatively unaffected by *real-time* top-down effects of arousal and attention are unsurprising given earlier observations that drugs that modulate arousal, or anesthesia do not affect them either (Kuwada et al., 2002; Cohen & Britt, 1982; Thornton et al., 1989). Thus, we conclude that these variables are not a significant factor except for cases where the signal is close to the noise floor where changes in measurement noise level can be consequential. Note that this discussion is strictly about endogenous (e.g., corticofugal) dynamic top-down effects on ABRs and EFRs. Sound-evoked efferent feedback effects could still contribute to variability in these measures, but are not discussed here. Also not discussed here are experience-dependent long-term plasticity effects that are thought to modulate subcortical responses (e.g., Kraus & Chandrasekaran, 2010).

## 4. Relationship between Candidate Measures

Next, we compare ABR, EFR and MEMR data across individuals to assess if they exhibit interrelationships consistent with cochlear synaptopathy when we incorporate some of the strategies discussed in Section 3. In particular, we use (1) the wideband MEMR measure (Keefe et al., 2017) with an FPL-calibrated 3-8 kHz broadband noise elicitor designed to produce a flat excitation pattern based on forward-masking based tuning estimates, (2) use the EFRs in response to a 3-8 kHz carrier noise band modulated at 223 Hz and with two different modulation depths (Bharadwaj et al., 2015), and (3) use the ABR wave I-to-wave V ratio, but using clicks restricted to the 3-8 kHz band. For comparing the ABR and the EFR, data were recorded at Boston University (N=30 ears, age 23-52 years, 12 female). For comparing the ABR and the MEMR, a different cohort of subjects (N=69 ears, age 18-50 years, 34 female) were recorded at Purdue University. All subjects had thresholds of 25 dB HL or better up to 8 kHz. Unfortunately, the data collection at Boston University was done before we started routinely acquiring audiometric data in the 9-16 kHz range; however, extended high-frequency audiograms were available for the 69 ears measured at Purdue University. As shown in Figures 6A and 6B, the ABR-EFR pair of measures, and the ABR-MEMR pair of measures each exhibit significant across-subject correlations, indicating contributions from a common underlying physiological factor. The ABR vs. MEMR correlations were significant even after adjusting for the variations in high-frequency (9-16 kHz) thresholds. This adjustment is conservative, as described in section 3.1. We interpret these observations as showing that there are individual variations consistent with variations in the degree of cochlear synaptopathy across these listeners. To test whether that interpretation in indeed the right one, future work will compare these measures on individuals particularly at risk for synaptopathy, either by virtue of their above-average acoustic exposures, or their age.

**Figure 6.**
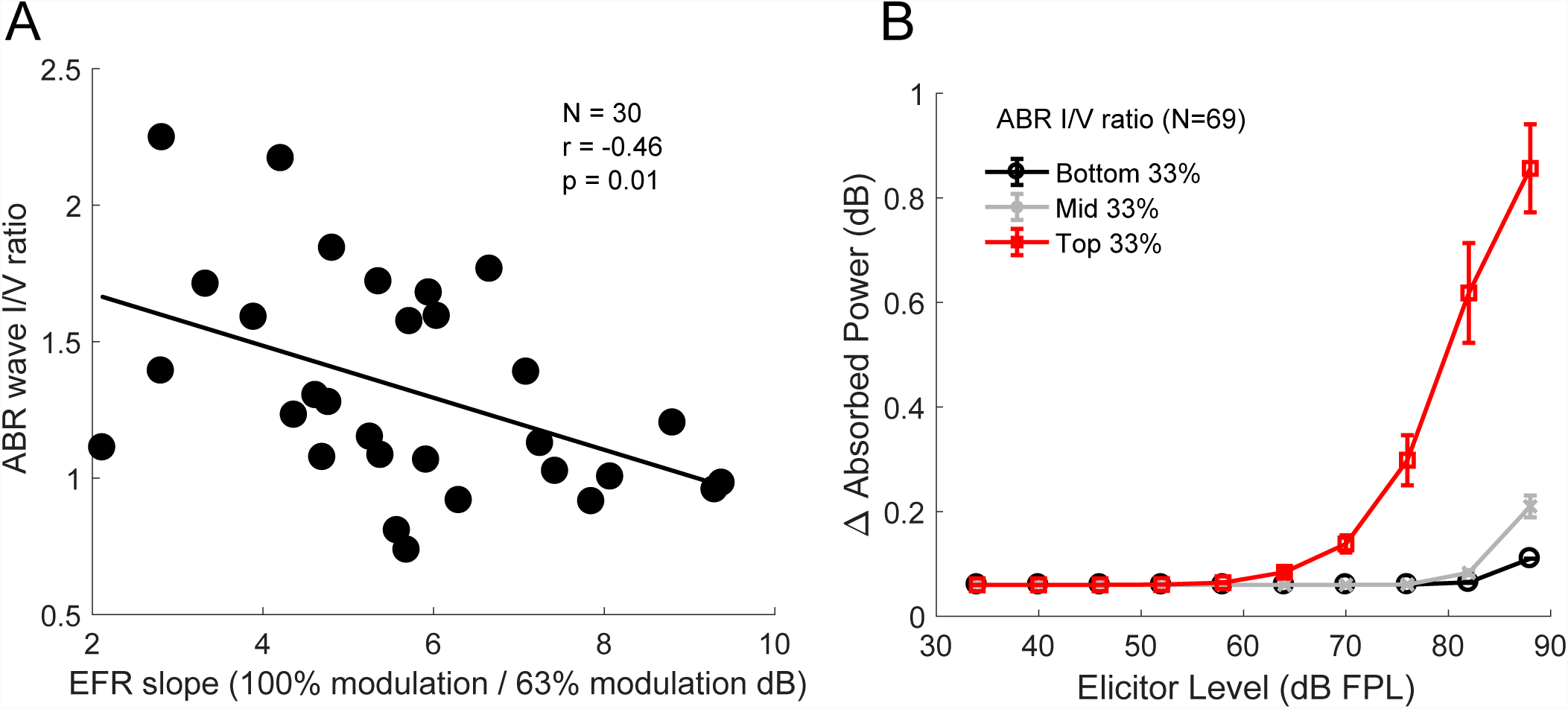
Across-individual correlations between the ABR (wave I/V ratio), EFR (change with modulation depth) and the MEMR (wideband average). Note that the eliciting stimulus for all three measures was restricted to the 3-8 kHz band, the region where “noise notches” often appear in human audiograms. Panel A shows the relationship between the ABR and the EFR for 30 ears with normal audiometric thresholds up to 8 kHz. A steeper EFR reduction with drop in modulation depth is associated with a smaller wave I/wave V ratio. Panel B shows the relationship between the ABR wave I/V ratio and MEMR measures from 69 ears with normal audiometric thresholds up to 8 kHz. In panel B, in order to show the association between the ABR and entire MEMR growth function, subjects were split into three groups based on their ABR wave I amplitudes (rather than show individual data points, which complicated visualizing the growth function). The median MEMR curve is shown for each group with error bars showing the.standard-error of the mean. Larger wave I/V ratios are associated with lower MEMR thresholds, and larger suprathreshold MEMR amplitudes. These observations are consistent with cochlear synaptopathy being a common source of individual differences in these measures.

## 5. Discussion

The robust finding of cochlear synaptopathy in multiple mammalian species raised the question of whether humans also exhibit synaptopathy (especially noise-induced synaptopathy), whether it may be measurable non-invasively in humans, and whether there are perceptual consequences to such damage. Unfortunately, assays that reliably reflect cochlear synaptopathy in specific strains of mice are affected by several extraneous factors in humans and other genetically hetogenous cohorts of animals. Here, through data from illustrative experiments, we discussed six such extraneous factors that could affect ABR, MEMR, and EFR measures. These experiments help us understand their effects and motivate strategies that may help mitigate them. While factors such as cochlear mechanical dispersion, audiometric loss at extended high frequencies, anatomical factors, and the stereotypical spectral response profile for the MEMR may be individual specific (and hence repeatable in a given individual), they nonetheless can obscure the effects of cochlear synaptopathy. Thus, a high degree of test-retest reliability by itself is insufficient for a candidate assay. The true test of whether a measure is potentially a good assay is whether the measure can capture individual variations in synaptopathy over and beyond the variance that is imposed by the host of extraneous variables. Indeed, by using methods that should mitigate the effects of some of these extraneous variables, we showed that the ABR wave I/wave V ratio for high-pass clicks, the wideband MEMR elicited by FPL-calibrated high-pass noise, and the modulation depth-dependence of the EFR elicited by modulated high-pass noise exhibit correlations with each other. This raises the possibility that cochlear synaptopathy might indeed be a widespread occurrence in humans—even those with normal hearing thresholds in ranges tested by typical audiometric screenings— and that the variations in the degree of synaptopathy might be the common factor resulting in correlations between these measures. Whether this is the case or not should be carefully explored in future studies. One line of investigation that would be particularly useful is to study these candidate non-invasive assays in genetically heterogeneous groups of animals where synaptopathy can be directly assayed using immunolabeling, and then comparing these metrics to the degree of synaptopathy observed.

For understanding of the prevalence and consequences of cochlear synaptopathy in humans, it is useful to separately consider those two aspects of the question, i.e., (1) Does synaptopathy (especially noise-induced) occur in humans, just as in rodents? (2) does it have perceptual consequences? There are many remaining barriers that complicate our ability to comprehensively answer these two questions, as illustrated in Figure 7. In behaving humans, risk factors for synaptopathy have to be estimated and then compared to some measured outcome/effect. There are many sources of variability in both arms of such experiments. While chronological age is easy to quantify, noise exposure history is not. One approach to reduce the estimation variability of noise-exposure risk is to study a group of individuals who are regularly and substantially overexposed compared to the average person (e.g., comparing occupationally exposed individuals to random age-matched individuals). Even if age and noise-exposure risk factors are well estimated, it is likely that individuals vary in their susceptibility to these risks (Davis et al., 2003; Maison et al., 2013; Liberman et al., 2014). Currently, it is unknown whether individual variations in susceptibility are large enough to overwhelm the effects of exposure, per se. This is analogous to the difficulty faced in relating individual outcome measures such as height (or) weight of children to diet – while it is generally accepted now that diet influences both height and weight, this was difficult to definitively establish given that heritability of human height is about 80% (Silventoinen et al., 2003).

**Figure 7.**
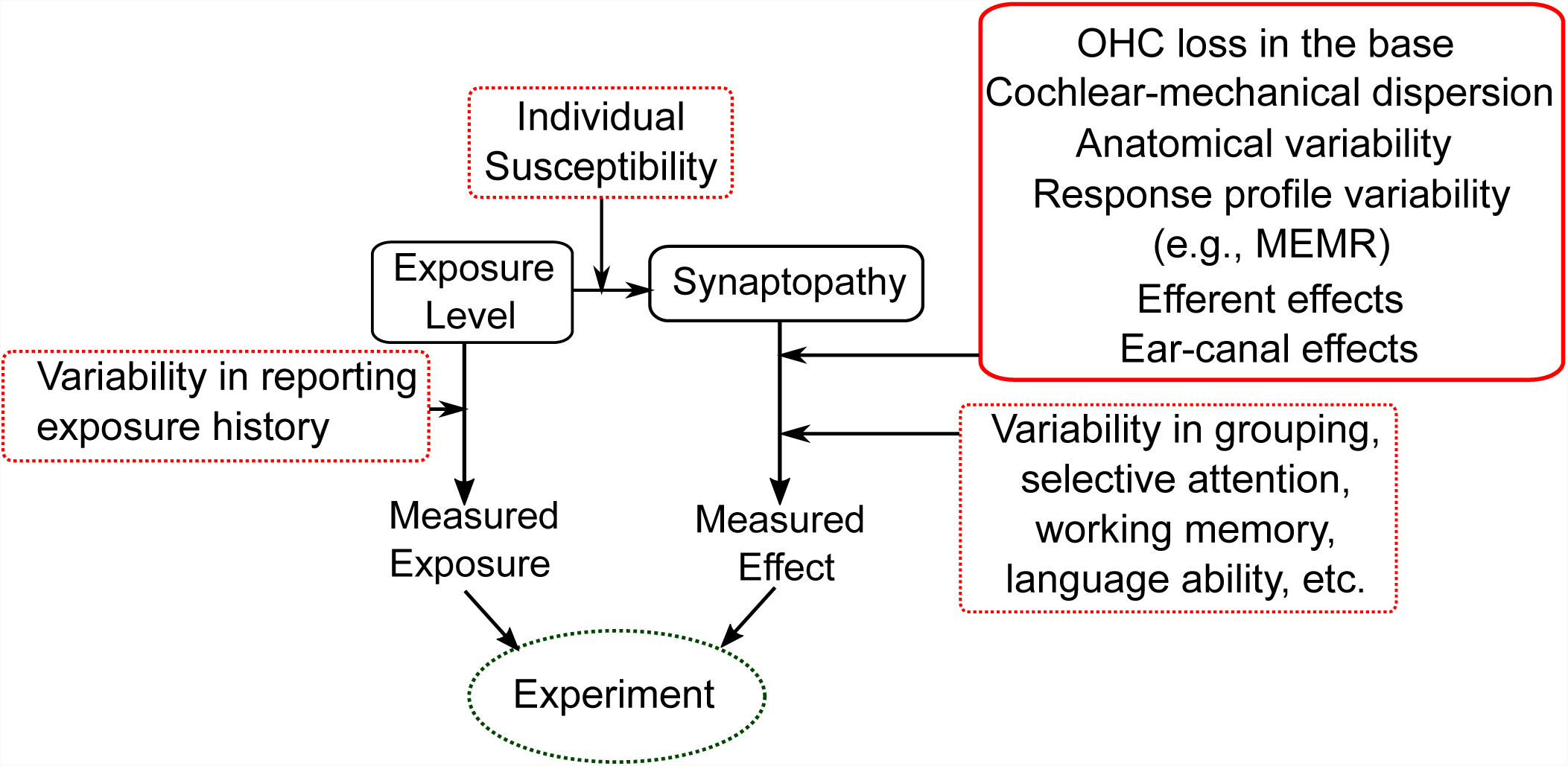
A schematic illustration of the challenges in establishing the prevalence and perceptual consequences of cochlear synaptopathy in humans. There are many sources of variability (illustrated in red boxes) in estimating both individual risk for synaptopathy and in the outcome measures available. For physiological assays, many of the factors illustrated in this manuscript can contribute to variability (solid red box). For perceptual outcome measures, still more factors could obscure the relationship between synaptopathy and perception (dashed red boxes). These sources of variability present a significant challenge for future studies in humans.

On the outcome-measure side of experiments exploring the effects of synaptopathy, many factors besides synaptopathy itself can affect our measurements. Here, it is useful to consider physiological assays of outcome, as considered in this report, or perceptual outcome measures. As discussed in Section 3, many factors can affect even physiological measures that originate early along the auditory pathway (also illustrated in Figure 7). On the other hand, when considering perceptual outcomes, even more factors become important. Firstly, in designing perceptual experiments, the neural code for many aspects of perception are unknown (although some aspects of physiology can be reasonably thought to underlie certain perceptual abilities, e.g., ITD processing in the MSO). Moreover, the effects of synaptopathy by itself may be small in simple perceptual tasks (Oxenham, 2016). It seems likely that perceptual effects of synaptopathy become apparent only in complex tasks such as speech identification in considerably adverse backgrounds, which rely on robust encoding of spectro-temporal variations through time in sound that is typically comfortably loud (and where the most vulnerable higher-threshold auditory nerve fibers are perhaps relatively more important for coding). If that were the case, one big challenge facing us is that of identifying task conditions where performance is truly limited by early sensory factors. Indeed, one recent large study that compared raw ABR amplitudes to speech-in-noise performance did not find them to be correlated (Smith et al., 2018). This is consistent with the notion that typical speech identification-in-noise task performance may be limited by “informational masking” (Brungart et al., 2006). Some studies that did find an association between early neural responses and selective attention tasks have used strategies that make it more likely that performance variations are limited by early sensory factors (Bharadwaj et al., 2015; Ruggles & Shinn-Cunningham, 2011). These strategies include (1) matching the target and masking sounds in monotone pitch so that the listener has to rely on subtle spatial cues to perform the task, (2) high-pass filtering of speech tokens so that the coding of envelopes of 3-8 kHz carriers become more important, and (3) using reverberation to degrade the temporal cues further exaggerating the importance of high-fidelity peripheral coding. Such strategies may be important because, although cochlear synaptopathy has received attention as one potential cause of degraded speech-in-noise perception (Plack et al., 2016; Bharadwaj et al., 2014; Liberman et al. 2016), it is just one factor that could contribute to outcomes. Successful listening in complex conditions not only relies on reliable coding of information at the auditory periphery but also successful scene segregation, selective attention, and other higher-level cognitive factors (Shinn-Cunningham, 2008). It is currently unclear whether the deficits in commonly-used speech-in-noise tasks are due to cochlear synaptopathy or problems with higher-level functions, or both. Evidence exists of individual differences in auditory grouping (Teki et al., 2013), selective attention (Choi et al., 2014; Bressler et al., 2014), working memory, and mapping of the target speech to articulatory sequences and meaning (Du et al., 2014; Wong et al., 2009). Because of the large number of variables that ultimately determine complex task performance, it may very well be the case that in order to establish any one of these factors as a contributor, we may have to study many of them in conjunction. One approach is to use candidate assays of as many of these variables as possible and study a large cohort of individuals while modeling their performance as dependent on both subcortical and cortical markers of the different variables that affect performance.

Given the many factors contributing to both our estimates of risk for synaptopathy, and outcomes thereof, it is perhaps prudent to interpret both positive and null association results in human experiments with caution. The experiments and considerations outlined here can help understand and reduce some of the sources of variability that affect three leading candidate physiological assays for cochlear synaptopathy. Here, we considered suprathreshold ABR wave I amplitudes and I/V amplitude ratios, EFR “slopes”, and the MEMR. Of these, the MEMR is perhaps the most promising candidate for a diagnostic measure of synaptopathy, both by virtue of how quickly it can be measured and owing to the possibility that it may particularly depend on higher-threshold auditory nerve afferents that are thought to be most vulnerable to damage. Notably, one measurement manipulation that we did not consider in this report is that of noise masking. It has been suggested that masked ABR and EFR measures may be useful in the diagnosis of cochlear synaptopathy by virtue of relative robustness of higher threshold low-spontaneous rate nerve fibers to masking (Mehraei et al., 2016; Paul et al., 2017; Bharadwaj et al., 2014). Future experiments should consider the use of these assays.

## Acknowledgements

This work was supported by NIH grants R01 DC015989 (HMB), R01 DC013825 (BGSC), and two emerging research grants (ERG) from Hearing Health Foundation (HMB and IC). We would like to thank Brooke Flesher, Kelsey Dougherty, and Anna Hagedorn for assistance with data collection at Purdue University, Karolina Charaziak for input during the initial stages of setting up the FPL calibration for the ER-10X system, and M. Charles Liberman for pointing us to relevant literature for Section 3.1.

